# Free Energy Principle in Human Postural Control System: Skin Stretch Feedback Reduces the Entropy

**DOI:** 10.1101/774026

**Authors:** Pilwon Hur, Yi-Tsen Pan, Christian DeBuys

## Abstract

Human upright standing involves an integration of multiple sensory inputs such as vision, vestibular and somatosensory systems. It has been known that sensory deficits worsen the standing balance. However, how the modulation of sensory information contributes to postural stabilization still remains an open question for researchers. The purpose of this work was to formulate the human standing postural control system in the framework of the free-energy principle, and to investigate the efficacy of the skin stretch feedback in enhancing the human standing balance. Previously, we have shown that sensory augmentation by skin stretch feedback at the fingertip could modulate the standing balance of the people with simulated sensory deficits. In this study, subjects underwent ten 30-second trials of quiet standing balance with and without skin stretch feedback. Visual and vestibular sensory deficits were simulated by having each subject close their eyes and tilt their head back. We found that sensory augmentation by velocity-based skin stretch feedback at the fingertip reduced the entropy of the standing postural sway of the people with simulated sensory deficits. This result aligns with the framework of the free energy principle which states that a self-organizing biological system at its equilibrium state tries to minimize its free energy either by updating the internal state or by correcting body movement with appropriate actions. The velocity-based skin stretch feedback at the fingertip may increase the signal-to-noise ratio of the sensory signals, which in turn enhances the accuracy of the internal states in the central nervous system. With more accurate internal states, the human postural control system can further adjust the standing posture to minimize the entropy, and thus the free energy.

## Introduction

Upright standing is one of the most important motor tasks that enable the use of the upper limbs and hands for tool use and dexterity. Even though this motor task seems trivial, it involves the complicated interplay between neurophysiology and biomechanics of sensory, neuromuscular, and neural processing subsystems^1–8^. Specifically, upright standing involves the integration and transformation of various sensory inputs (e.g., vision, vestibular system, proprioception, touch) via a highly-coordinated central nervous system (CNS) into bodily information (e.g., position, velocity, and acceleration of body sway). Degradation in the sensory systems (e.g., vision^7, 9^ proprioception^10–12^, vestibular system^8, 13^, touch^3, 14^), muscular systems^15, 16^ and nervous systems^17, 18^ worsens balance of the upright standing.

One of the key elements for the successful balance rehabilitation is to quantitatively assess the balance. Traditionally, descriptive parameters have been used to characterize the postural sway of the standing balance^19^. Several mathematical models using statistical mechanics have also been introduced to capture the stochastic behaviors of the postural sway^20–22^. However, these measures provide only the statistical description of the system, but not the dynamics itself^23^. Instead, several researchers tried to model the sensorimotor system from the control-theoretic perspectives to understand the human motor control system. Wolpert et al.^24^ proposed that a sensorimotor system has an internal model such that the CNS internally simulates the dynamical behavior of the system for the purpose of planning, control and learning. They argued that the CNS internally integrates sensory signals such that predicted sensory errors temporally propagate in the framework of a Kalman filter. In addition, motor learning was suggested to be the process of updating the prior knowledge of the system by evidences from the error feedback in the Bayesian-optimal framework^25^. However, even though the control-theoretic framework seems fascinating, it requires for the biological systems to have the complicated inverse model and the heavy computation, which are not likely to be realizable neurobiologically^26^.

Recently, a unifying theory, called free-energy principle, was proposed to account for the action, perception and learning of sensorimotor systems^27^. The free-energy principle argues that a self-organizing biological system at its equilibrium point tries to minimize its free energy. In other words, a self-organizing biological system tries to resist a tendency toward disorder. According to the free-energy principle, a self-organizing biological system minimizes the free energy either by i) updating the internal state (equivalently, internal model or prior^25^) or ii) correcting body movement with appropriate actions. Free energy principle does not require any complicated inverse models and is neurobiologically realizable^26^. Therefore, the idea of the free-energy principle may also be applied to human postural control system to understand how the human CNS maintains the standing balance under external perturbations. In other words, this framework can be used to quantify the human postural control system^7, 28^. In the Method section and Appendix, we will examine the use of the free-energy principle to derive metrics that quantify or assess the human standing balance.

Rehabilitation is a process of a motor adaptation in response to external changes or stimuli to restore the normative motor functions. These external changes or stimuli mostly result from errors or deviations from the reference (or desired) trajectories for the given tasks. However, how the motor adaptation takes place is still not well known and is one of the important research topics in human motor control and rehabilitation. One widely-accepted idea is that human sensorimotor systems utilize the error feedbacks for motor adaptation^29^. Several studies have reported that amplifying the movement errors could enhance the rehabilitation outcomes, which is sometimes referred to as an *error augmentation*^13, 30, 31^. Although rehabilitation may include internal sources of adaptation such as motor imagery, these are not the focus of this study.

Cutaneous feedback has been of a great interest as a way to sensory augmentation for the balance enhancement^13, 32–34^. Random vibrotactile feedback has been used to strengthen the weak signal, known as stochastic resonance, and thus improve the sensorimotor functions^35–38^. It was shown that the random vibrotactile signal at various body parts (e.g., foot sole, wrist) enhanced sensorimotor functions (e.g., light touch sensation, gait variability, hand functions) in young adults, elderly adults, and stroke patients, suggesting the great potential of the random vibrotactile feedback for balance rehabilitation^35, 36, 38–40^.

Similarly, Jeka et al.^32^ reported the effect of light touch in standing balance. They found that subjects who are lightly touching a fixed surface with their index fingertip (with the normal force less than 1N) exhibited the reduced postural sway in tandem Romberg posture. Clapp and Wing^33^ also showed a reduction in postural sway in the sagittal plane when subjects were making a light contact with their fingertips on a fixed surface during normal bipedal stance. Pan et al.^13^ showed that skin stretch and skin deformation at the fingertip could provide subjects with additional information to identify the direction of body sway and thus to modulate the postural sway. These suggest that additional cutaneous sensory inputs at the fingertip can provide a robust regulation of the postural sway. It has been shown, by partially blocking sensory afferents during human standing, that the enhanced balance due to light touch was not due to the mechanical support on the fingertip but due to the tactile feedback^41^. Krishnamoorthy et al.^34^ observed that a light touch at the head or neck reduced the body sway even more effectively compared with a fingertip touch.

The objective of this study is twofold: i) to formulate the human standing postural control system in the framework of the free-energy principle, and thus to introduce measures that quantify the human postural control system and ii) to investigate the efficacy of the skin stretch feedback in enhancing the human standing balance. We hypothesized that skin stretch feedback would both reduce the entropy of the postural sway and improve balance with respect to other measures in accordance with the free energy principle. The formulation of the human postural control system in the framework of the free-energy principle is given in the Method section. We also provide a proof that the long-term average of the free energy can be approximated by an entropy of the postural sway in the Appendix. In the Experiment section, the details of experimental protocol to test the efficacy of the skin stretch feedback are described. In the Result and Discussion sections, we show that the skin stretch feedback at the fingertip reduces Entropy of the postural sway.

## Methods

In this section, we introduce the framework of free-energy principle and provide an algorithm to estimate the entropy of the human postural sway. To test the feasibility of the framework, we conduct an experiment of quiet standing with and without skin stretch feedback.

### Free energy principle

The free-energy principle states that a self-organizing biological system at its equilibrium point tries to minimize its free energy^27^. Similar to the idea of the conventional optimal control theory based on forward-inverse models^24, 42, 43^, free energy principle explains how a biological system makes movements, perceives the environments and infers the internal states. However, unlike the conventional optimal control theory, free energy principle does not require the complicated inversion process of the forward models via heavy computation and optimization^26^. That is, free energy principle uses a forward model only and does not need to solve the complicated Bellman equation from dynamic programming with cost functions. Instead, it uses prior beliefs about the given tasks learned via previous sensorimotor behaviors, which is generally considered to be Bayes-optimal^44, 45^. With these benefits, we frame the human postural control system with the free energy principle.

#### Surprise

In essence, free energy principle, or equivalently active inference, claims to minimize the prediction error of the forward model (i.e., generative model) via appropriate actions and perceptions^27^. The prediction error is called self-information, or *surprise*. In other words, minimizing the surprise is equivalently maximizing the accuracy of prediction about the sensory states by the agent (i.e., biological systems). A careful mathematical formulation (see Appendix) can prove that minimizing the free energy is implicitly equivalent to minimizing surprise. To minimize the free energy, the agent either i) makes a movement of the body to alter the sensory input or ii) updates the internal states of the brain to refine the approximation of the posterior beliefs (i.e., recognition model)^27^. When this principle is applied to the human standing balance, CNS is understood to minimize the free energy either by correcting body posture or by updating the internal model of the postural control system (i.e., self-awareness of body movement). In other words, the human postural control system with the goal or intention of stable standing balance always tries to be confident about the the current state of its posture.

#### Entropy

The long-term average of surprise can be proven to be *Entropy* (see Appendix). Therefore, the free energy principle is called a *minimum entropy principle*, suggesting that CNS keeps the average surprise of the body posture sampled from a probability distribution or density to be low. A density with low entropy means that, on average, the body posture is relatively predictable^23, 27^.

Then, what is the meaning of entropy minimization in the problem of the human postural control? Ones are easily tempted to consider that the postural control system is trying to prevent any postural sways. From the optimal control point of view, this may seem to be true since the usual cost functions include states (e.g., linear quadratic regulator). However, it is important to note that the equilibrium point during the quiet standing is not necessarily the same as standing still with no postural sway. In general, humans do not take the strategy of standing still since it requires excessive efforts^46^. Instead, the human postural control system allows flexibility to some extent such that whenever the postural sway is within a certain range, little control efforts are made in correcting the posture^20^.

In our previous study^23^, we developed an algorithm to compute the entropy of human quiet standing and showed that people with better balance control have smaller entropy of postural sway distribution. Postural sway data captured by the center of pressure (COP) were used to extract the reduced order dynamics of the human postural control system in the framework of a Markov chain. The reduced order dynamics were expressed in the form of transition probability matrix that describes the movement of COP, and thus the evolution of COP distribution. With the careful development of the algorithm, the distribution is guaranteed to converge to be stationary. This framework, a free energy principle applied to human postural control system, is called *Invariant Density Analysis* (see Data analysis section). For the detailed development of the framework, please refer to Hur et al.^23^. From the *Invariant Density Analysis*, it was found that people with degraded sensorimotor functions or under perturbations had significantly higher entropy compared with healthy people with no perturbations^16, 23^.

### Skin stretch feedback

#### Reduction of entropy

As mentioned in the Introduction, skin stretch feedback showed its potential for balance rehabilitation. Skin stretch is a simple mechanism for generating additional sensory signals and can be made accessible in almost every situation via fixed structures or wearable robotic devices. Skin stretch feedback is a type of *sensory augmentation* that enables the *error augmentation* for motor learning. In terms of free energy principle, the additionally-sampled sensory data (i.e., skin stretch feedback) increase the signal-to-noise ratio (SNR), and thus the brain can generate motor commands more effectively by maximizing the model evidence. In other words, the increased prediction accuracy of the sensory states due to skin stretch feedback, and thus the increased SNR contributes to the optimal motor planning. The motor adjustment from the optimal motor planning further contributes to minimizing the prediction errors. These repeated processes eventually minimize the entropy of sensory state probability^27^.

#### Skin stretch feedback device

Previously, we developed a wearable sensory augmentation system for standing balance rehabilitation using a skin stretch feedback device (Fig. 1)^13^. The device was inspired by the concept of a light touch as mentioned in the Introduction. By mimicking the shear force that subjects may experience at the fingertip during a light touch of fixed surface, the additional postural sway information was provided to the subjects’ fingertip in the form of the skin stretch feedback. This preliminary study^13^ found that the sensory augmentation due to skin stretch feedback at the fingertip could enhance the standing balance, indicating the reinforcement of the accuracy of the sensory prediction.

**Figure 1.**
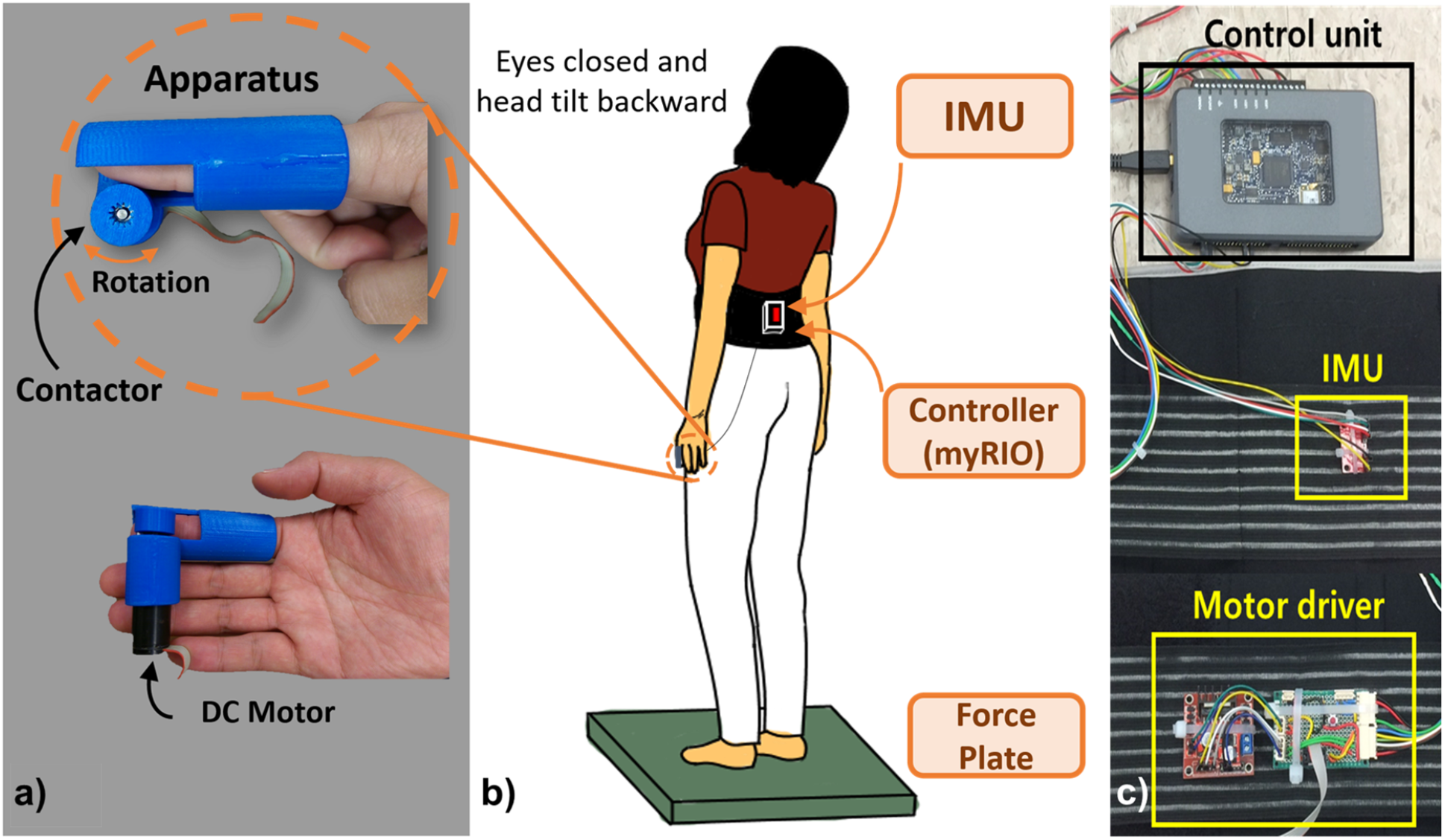
Skin stretch feedback system and experiment setup. The system consists of a skin stretch device (SSD), a waist belt enclosing an inertia measurement unit (IMU), a motor driver, and an embedded control unit (myRIO). a) SSD is designed to be attached to subject’s index finger. A DC motor is mounted inside the housing of SSD. A contactor rotates while the subject is swaying back and forth to provide additional cutaneous skin stretch feedback on one’s fingertip pad. b) Subjects were asked to perform 30 seconds of quiet standing on a force plate with simulated sensory deficits (eyes closed and head tilted backward).

The components of the system include (Fig. 1): a wearable skin stretch device (SSD), and a waist belt enclosing an inertia measurement unit (IMU) (MPU-9150, InvenSense Inc., San Jose, CA), a motor driver (L298N, STMicroelectronics, Italy), and an embedded control unit (myRIO, National Instruments, Austin, TX). A wearable skin stretch device (Fig. 1a) is designed to be attached to the subject’s index finger. SSD roughly imposed about 20-30g weight on the fingertip^13^ and the coefficient of friction between the contactor and the fingertip was roughly 0.5 or less^47^. A DC motor (1524T009SR, Faulhaber, Germany) mounted inside the device’s housing was powered by an h-bridge type motor driver (Fig. 1c). As the subject sways, IMU detects the change of the body orientation and the motor rotates accordingly. Then, the contactor stretches the fingertip pad, which induces the skin stretch feedback. The housing of the device was made of ABS-type filament through a 3D printer (Replicator 2X, Makerbot, Brooklyn, NY). Different sizes of housings and contactors were fabricated to cover various finger sizes of all subjects so that the contact at the fingertip was maintained with qualitatively small normal force.

#### Control strategy

Subjects wore the SSD at their index finger and the waist belt around their waist. The IMU was placed at the estimated center of mass (COM) location (Fig. 1b). Postural sway in anterior-posterior (AP) direction was registered by the IMU (e.g., pitch angle). The recorded sway angle was processed by myRIO in real-time. The contactor of SSD was controlled by proportional, integral and derivative (PID) terms of body pitch angle, *θ*_*p*_. Specifically, body pitch angle was mapped to the angular velocity of the contactor, *ω*_*c*_, of SSD as follows:

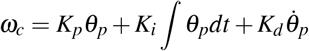

where *K*_*p*_, *K*_*i*_, and *K*_*d*_ are PID gains with units of *s*^−1^, *s*, and unitless, respectively. To find *K_d_*, we tuned the parameters during the pilot test to make sure the subjects do not feel any discomfort, but feel *light touch* at their fingertips. We let *K*_*p*_ = *K*_*i*_ = 0 since sway velocity information was found to be a more accurate form of information acquired by sensory systems when compared to its sway position and acceleration information for human standing balance^5, 48^. Intuitively, this idea of *D* control is plausible since skin stretch feedback is based on the flow or slip at the contact surface. To avoid any potential discomfort at the fingertip, the maximum angular velocity of the contactor was limited to 25 *rad/s*.

Both direction and velocity information of postural sway can be provided and augmented by additional light touch of a stationary surface^48–51^. Moreover, it was found that postural sway was highly consistent with the driving frequency of moving surface where subjects are touching. This emphasizes the important role of cutaneous feedback in modulating the control of upright posture^50, 51^. The maximum value of *ω*_*c*_ was restricted to 25 rad/s for this purpose, which is about 4 Hz. In this way, when the subject stayed still regardless of the pitch angle (e.g., forward or backward), the contactor stopped, and no skin stretch was applied to the fingertip pad. This also removed the drift issues of the IMU data.

### Experiment

To test the idea of free energy principle and the efficacy of the skin stretch feedback in reducing the entropy of the standing balance, the following experiment was conducted.

#### Subjects

In Pan et al.^13^, the range of COP in AP direction with two factors (sensory augmentation and vision) had the Partial *η*^2^ = 0.1. We assumed a correlation of 0.5 between these two data; then we could get an effect size of 0.33. Then, at least 12 subjects were needed for each group for a significance level of 0.05 and a power of 0.95 using G*Power^52^. To ensure significance, 15 subjects were recruited (three females and twelve males: mean age s.d.: 25.6 ± 3.33 years) for this experiment. The subjects have neither been neurologically impaired nor had balance issues before. They were informed of the experimental procedures except how the sensory feedback device operates. All subjects provided the informed consent before the experiment started. All experimental protocol and methods in this experiment were performed in accordance with the relevant guidelines and regulations from the Declaration of Helsinki and were approved by the Texas A&M University Institutional Review Board.

#### Protocols

Subjects wore the SSD on their index finger and put on the waist belt around their waist. Their arms hung naturally by their sides. Ten trials of 30 second quiet standing on a forceplate (OR6, AMTI, Watertown, MA) with simulated sensory deficits and two sensory augmentation conditions were performed. Subjects were instructed to stand quietly. No other instructions were given to subjects about the behavior of the contactor. To simulate the sensory deficit condition, both visual and vestibular deficits were induced by asking subjects to close their eyes and tilt their head up at least 45° (Fig. 1b). Under such a head-extended condition, the vestibular organ moves and the utricle otoliths are put into irregular location, which perturbs the vestibular sensory system and therefore results in postural imbalance^53, 54^. Subjects put on an overhead safety harness for protection against unwanted and unexpected falls. Two sensory augmentation condition were included: i) SSD is turned on (ON), and ii) SSD is turned off (OFF). The experiment comprised of practice and main sessions. Practice session aimed at familiarizing the subjects with the experiment setup. Subjects were asked to stand quietly with barefoot on a forceplate. There were three practice trials before the main session to make sure that subjects become familiar with the experiment. In the main session, subjects were instructed to perform the same quiet standing tasks as in the practice session under the 2 sensory augmentation conditions: i) SSD OFF, ii) SSD ON. Each condition was repeated 10 times to minimize random effects by taking the mean values of each metric. Note that the order of all 20 trials was fully randomized, and subjects were not informed of the sensory augmentation condition. Break was provided upon request, and a two-minute rest was provided between every 5 trials to avoid muscle fatigue. The whole experiment lasted about 30 minutes. Note that in both sensory augmentation conditions, subject wore the SSD all times even when no skin stretch feedback was provided.

#### Data analysis

Based on the free energy principle, human postural control system tries to minimize the entropy of body posture sampled from a postural sway probability density. An algorithm, Invariant Density Analysis (IDA), to compute the probability density of body posture was developed by Hur et al.^23^. In the following, a brief explanation of IDA is presented. IDA includes several measures including entropy to parameterize the invariant density. In addition to IDA, Stabilogram Diffusion Analysis (SDA) is computed to complement the explanation of the postural control.

Postural balance was quantified by measuring the COP displacement registered by a forceplate (OR6, AMTI, Watertown, MA, USA) and a data acquisition system (USB-6002, National Instruments, Austin, TX). COP from a forceplate and pitch angle from IMU were collected at a sampling rate of 1 kHz and 500 Hz, respectively. The data were processed offline using MATLAB (R2016a, MathWorks, Natick, MA). Postural sway measures were computed in anterior-posterior (AP), medio-lateral (ML), and radial (Rad) direction. Rad displacements are the Euclidean distance from the mean COP to each COP.

### Invariant density analysis

Previously, we^23^ introduced a reduced-order finite Markov chain model to analyze the stochastic structure of postural sway, which provides insight into the long-term system behavior. This model describes the states (zero-mean COP data) of the dynamical system and evolution of those states. It was found that the distribution of COP over the state space would eventually converge to a unique steady state distribution *π*, known as invariant density. To acknowledge the details of subject-specific COP behaviors, five parameters were introduced as follows. *P*_*peak*_ is the peak value of *π*. *D̄* is the average location (state) of the COP, and the unit is *mm*. *D*_95_ is the largest state at which there is a 95% probability of containing the COP. It has the same unit as *D̄*. *λ*_2_ is the second largest eigenvalue of the transition matrix and describes the convergence rate of the system to its invariant long-term behavior. *H* is the entropy and is the measure of randomness. *H* is computed as *H* = − ∑_*i*∈*I*_ *π*(*i*) log_2_ *π*(*i*) where *I* is the set of all possible states.

In addition to the above five parameters, we computed a metric that may provide insight into the control mechanisms of the postural control system. The eigenvector corresponding to the second largest eigenvalue is of our interests^28^. It was noted that investigation of the eigenvector corresponding to *λ*_2_ has the potential to provide a complete understanding of the embedded dynamics in the reduced order model^28^. For simplicity, we call the eigenvector corresponding to the second largest eigenvalue as the second eigenvector. Recently, the second eigenvector has been used to formulate an intuitive understanding of the dynamics for a finite state-space ergodic Markov chain by decomposing the state space into essential features^55–57^. Specifically, Dellnitz^55^ reported that the state space could be subdivided into two subsets depending on the sign of the corresponding second eigenvector (i.e., positive vs. negative). Each subset shows different dynamic behavior. Using this idea, we investigated the zero crossing point of the second eigenvector, 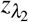.

### Stabilogram diffusion analysis

Collins and DeLuca^20^ proposed a Stabilogram Diffusion Analysis (SDA) that could provide several physiologically-meaningful parameters from stabilogram from the viewpoint of statistical mechanics. SDA computes how the COP diffuses over time at each arbitrary time. It was shown that the postural control system during quiet standing could be separated into two different control mechanisms: open-loop and closed-loop control. Figure 2 shows an example of a resultant planar stabilogram-diffusion plot. Short-term and long-term regions are dominated by the open-loop and closed-loop control strategies, respectively. This phenomenon can be described by the following parameters.

**Figure 2.**
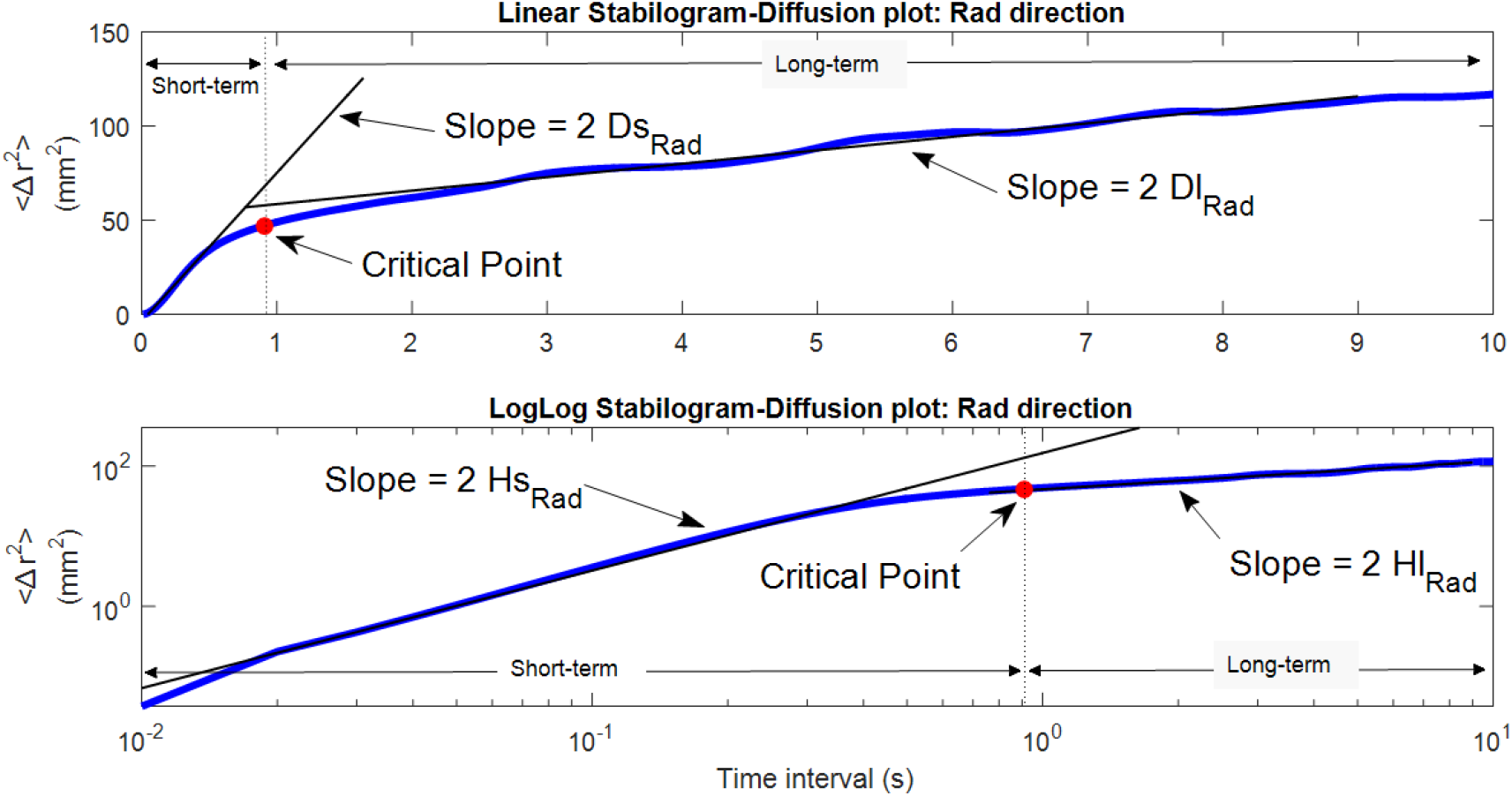
Linear and log-log stabilogram diffusion plots in the radial (Rad) direction. Short-term and long-term regions fitted by straight black regression lines are dominated by the open-loop and closed-loop control strategies respectively. Stabilogram diffusion parameters (*D*_*s*_, *D*_*l*_, *H*_*s*_, *H*_*l*_) are determined by the slopes of lines fitted to short-term or long-term regions. A critical point is defined as the point where the slope changes considerably from one region to another.

Diffusion coefficients (*D*_*s*_, *D*_*l*_), which equal one-half of the slopes of the fitting line to both regions, represent the level of stochastic activities of the COP about the plane of support. For example, greater *D*_*s*_, compared to *D*_*l*_, reflects a more random behavior and less active control in the short-term interval. The next parameters are called the scaling exponents (*H*_*s*_, *H*_*l*_), which are the slopes of the log-log plots of the stabilogram, evaluate the persistence/anti-persistence of the COP behavior. Scaling exponents suggested that COP tends to move away from the equilibrium during the short-term interval (*H*_*s*_ > 0.5, persistent) whereas COP tends to return to the equilibrium over long-term interval (*H*_*l*_ < 0.5, antipersistent). The last two parameters are critical point coordinates (Δ*t*_*c*_, ⟨Δ*r*^2^⟩_2_) where ⟨·⟩ represents the mean value. A critical point is defined as the point where the slope changes considerably from one region to another, representing the critical time interval (Δ*t*_*c*_) and critical mean square displacements (⟨Δ*r*^2^⟩_*c*_). The critical point is calculated by finding the first minimum of the second derivative of the linear stabilogram-diffusion plot.

#### Statistical analysis

Data from all trials were averaged for the two feedback conditions, SSD OFF and SSD ON. A two-tailed paired *t*-test was performed using SPSS statistical software package (v21, SPSS Inc., Chicago, IL) to evaluate the effects of SSD on quiet standing balance analyzed using three different methods. The significance level was set to *α* = 0.05.

## Results

Effect of the skin stretch feedback (SSD) on standing balance under the simulated sensory deficit condition is summarized in Table 1. Specifically, among IDA measures, *P*_*peak,AP*_ (*t*(14) = 2.12, *d* = 0.55, *p* = 0.05 where *t*, *d*, and *p* represent *t* statistic, Cohen’s *d* for effect size and *p* value) significantly increased, and *H*_*AP*_ (*t*(14) = −2.11, *d* = 0.54, *p* = 0.05), *H*_*Rad*_ (*t*(14) = −2.34, *d* = 0.60, *p* = 0.04), and 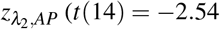, *d* = 0.66, *p* = 0.02) significantly decreased when SSD was provided. Even though no significant differences were found in the other IDA measures, *D*_95,*Rad*_ (*t*(14) = −1.94, *d* = 0.50, *p* = 0.07) and 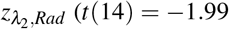, *d* = 0.51, *p* = 0.07) tended to decrease when SSD was provided. Figure 3 shows the representative invariant density plots of SSD ON and OFF conditions, and their associated IDA parameters (e.g., *P*_*peak*_, D̄, and *D*_95_). It is observed that when SSD was on, *P*_*peak,AP*_ increased, whereas *H*_*AP*_, and *H*_*Rad*_ significantly decreased. Also, *D*_95,*AP*_ tended to decrease when SSD was on. Figure 4 shows the representative plots of the second eigenvector of the transition matrix for both SSD ON and OFF conditions. It is observed that 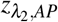 significantly reduced when SSD was on. Also, 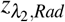 tended to reduce when SSD was provided.

**Table 1.**
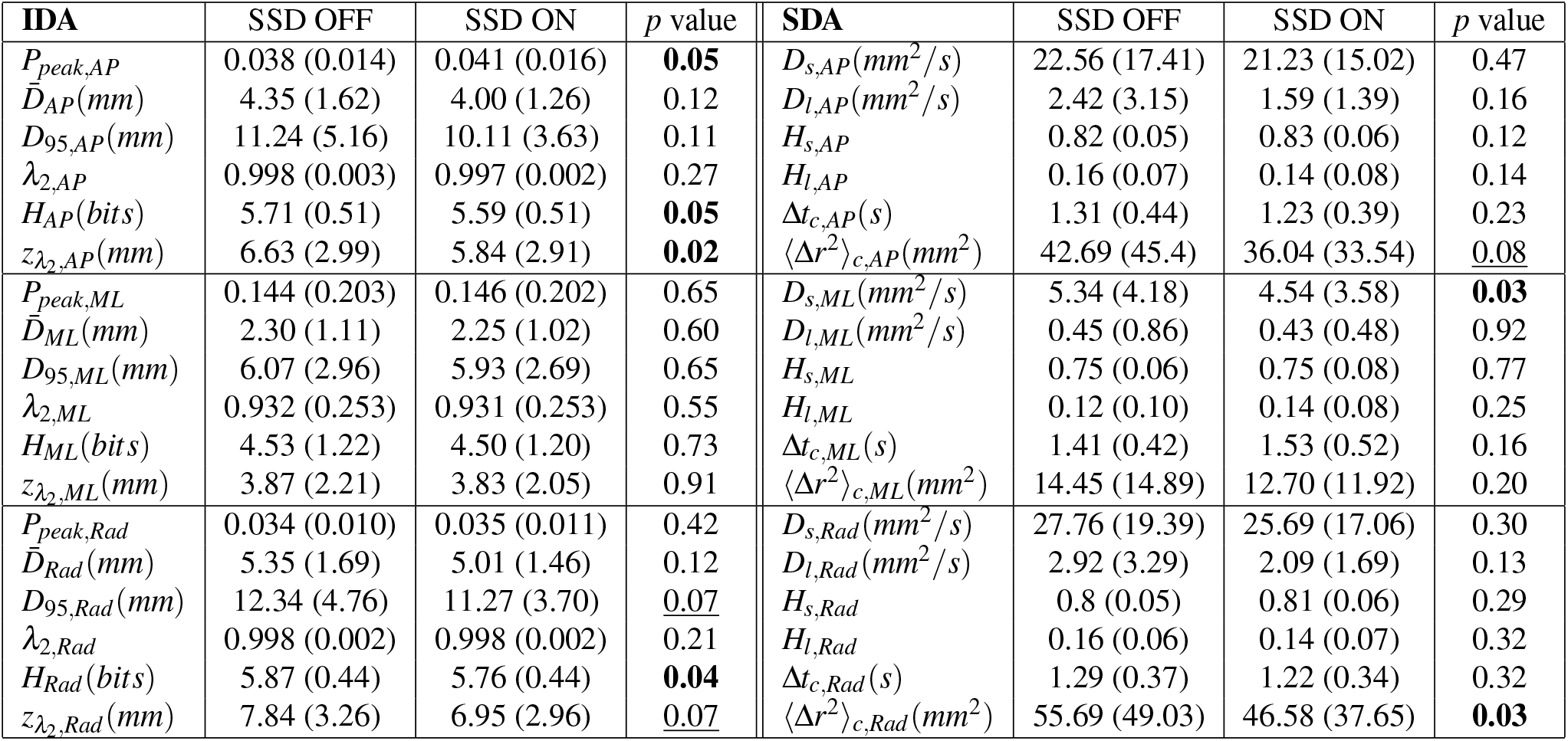
IDA and SDA measures of postural sway in AP, ML and Rad directions. Value represents mean (standard deviation) with the condition of skin stretch feedback (SSD). Bold text denotes significant differences between SSD OFF and SSD ON) (*p* < 0.05). Underline denotes near significance or tendency (*p* < 0.10).

| IDA | SSD OFF | SSD ON | $p$ value | SDA | SSD OFF | SSD ON | $p$ value |
| --- | --- | --- | --- | --- | --- | --- | --- |
| $P_{peak,AP}$ | 0.038 (0.014) | 0.041 (0.016) | **0.05** | $D_{s,AP}(mm^2/s)$ | 22.56 (17.41) | 21.23 (15.02) | 0.47 |
| $\bar{D}_{AP}(mm)$ | 4.35 (1.62) | 4.00 (1.26) | 0.12 | $D_{l,AP}(mm^2/s)$ | 2.42 (3.15) | 1.59 (1.39) | 0.16 |
| $D_{95,AP}(mm)$ | 11.24 (5.16) | 10.11 (3.63) | 0.11 | $H_{s,AP}$ | 0.82 (0.05) | 0.83 (0.06) | 0.12 |
| $\lambda_{2,AP}$ | 0.998 (0.003) | 0.997 (0.002) | 0.27 | $H_{l,AP}$ | 0.16 (0.07) | 0.14 (0.08) | 0.14 |
| $H_{AP}(bits)$ | 5.71 (0.51) | 5.59 (0.51) | **0.05** | $\Delta t_{c,AP}(s)$ | 1.31 (0.44) | 1.23 (0.39) | 0.23 |
| $z_{\lambda_2,AP}(mm)$ | 6.63 (2.99) | 5.84 (2.91) | **0.02** | $\langle \Delta r^2 \rangle_{c,AP}(mm^2)$ | 42.69 (45.4) | 36.04 (33.54) | <u>0.08</u> |
| $P_{peak,ML}$ | 0.144 (0.203) | 0.146 (0.202) | 0.65 | $D_{s,ML}(mm^2/s)$ | 5.34 (4.18) | 4.54 (3.58) | **0.03** |
| $\bar{D}_{ML}(mm)$ | 2.30 (1.11) | 2.25 (1.02) | 0.60 | $D_{l,ML}(mm^2/s)$ | 0.45 (0.86) | 0.43 (0.48) | 0.92 |
| $D_{95,ML}(mm)$ | 6.07 (2.96) | 5.93 (2.69) | 0.65 | $H_{s,ML}$ | 0.75 (0.06) | 0.75 (0.08) | 0.77 |
| $\lambda_{2,ML}$ | 0.932 (0.253) | 0.931 (0.253) | 0.55 | $H_{l,ML}$ | 0.12 (0.10) | 0.14 (0.08) | 0.25 |
| $H_{ML}(bits)$ | 4.53 (1.22) | 4.50 (1.20) | 0.73 | $\Delta t_{c,ML}(s)$ | 1.41 (0.42) | 1.53 (0.52) | 0.16 |
| $z_{\lambda_2,ML}(mm)$ | 3.87 (2.21) | 3.83 (2.05) | 0.91 | $\langle \Delta r^2 \rangle_{c,ML}(mm^2)$ | 14.45 (14.89) | 12.70 (11.92) | 0.20 |
| $P_{peak,Rad}$ | 0.034 (0.010) | 0.035 (0.011) | 0.42 | $D_{s,Rad}(mm^2/s)$ | 27.76 (19.39) | 25.69 (17.06) | 0.30 |
| $\bar{D}_{Rad}(mm)$ | 5.35 (1.69) | 5.01 (1.46) | 0.12 | $D_{l,Rad}(mm^2/s)$ | 2.92 (3.29) | 2.09 (1.69) | 0.13 |
| $D_{95,Rad}(mm)$ | 12.34 (4.76) | 11.27 (3.70) | <u>0.07</u> | $H_{s,Rad}$ | 0.8 (0.05) | 0.81 (0.06) | 0.29 |
| $\lambda_{2,Rad}$ | 0.998 (0.002) | 0.998 (0.002) | 0.21 | $H_{l,Rad}$ | 0.16 (0.06) | 0.14 (0.07) | 0.32 |
| $H_{Rad}(bits)$ | 5.87 (0.44) | 5.76 (0.44) | **0.04** | $\Delta t_{c,Rad}(s)$ | 1.29 (0.37) | 1.22 (0.34) | 0.32 |
| $z_{\lambda_2,Rad}(mm)$ | 7.84 (3.26) | 6.95 (2.96) | <u>0.07</u> | $\langle \Delta r^2 \rangle_{c,Rad}(mm^2)$ | 55.69 (49.03) | 46.58 (37.65) | **0.03** |

**Figure 3.**
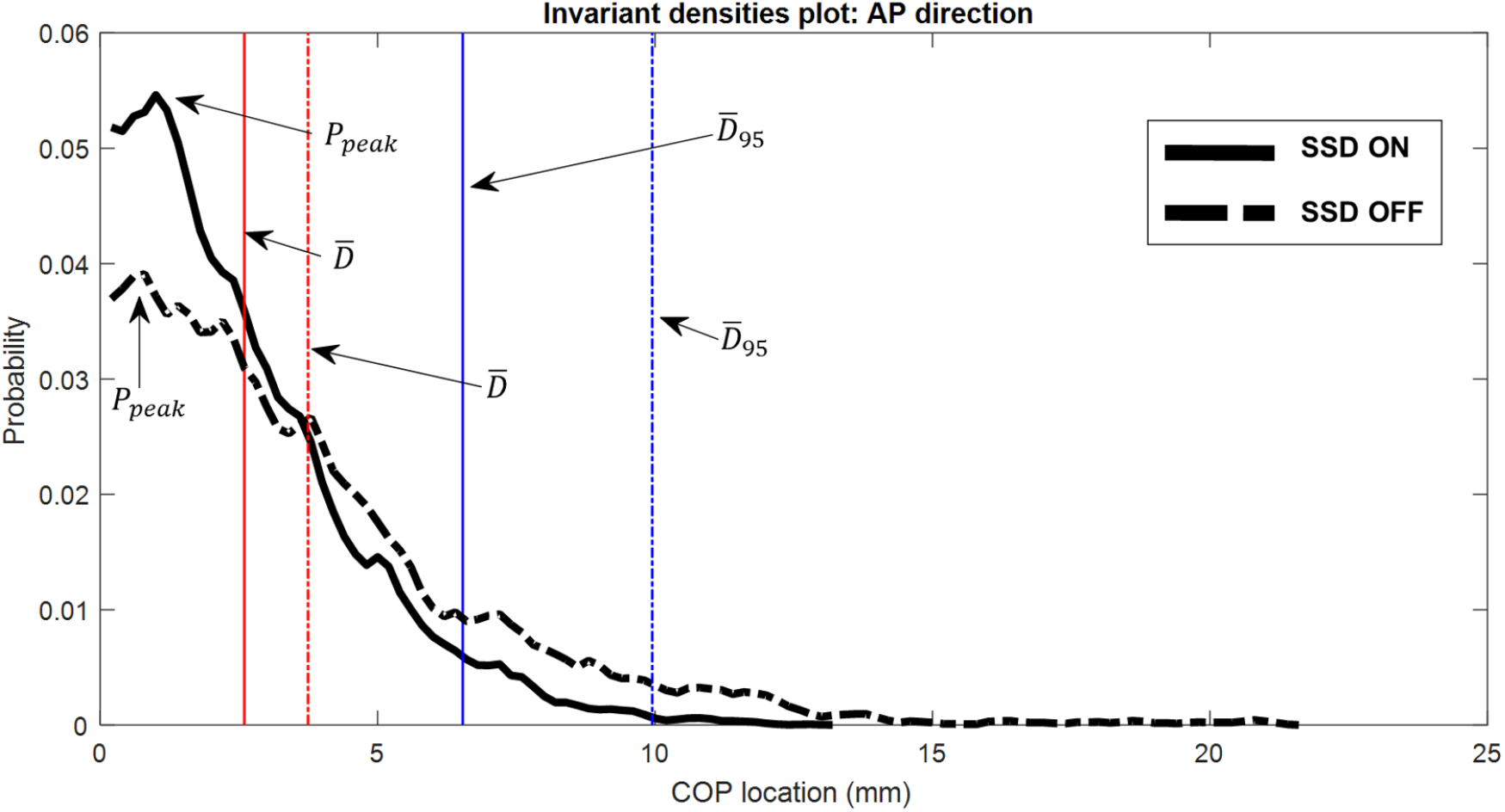
Representative IDA plots of both SSD ON (solid) and SSD OFF (dashed) conditions, and their associated IDA parameters. When additional skin stretch was applied on the subject (SSD ON), *P*_*peak*_ increased, whereas *D̄* and *D*_95_ decreased, indicating a higher probability that COP will be found closer to the equilibrium, and COP traveled within a smaller region near centroid.

**Figure 4.**
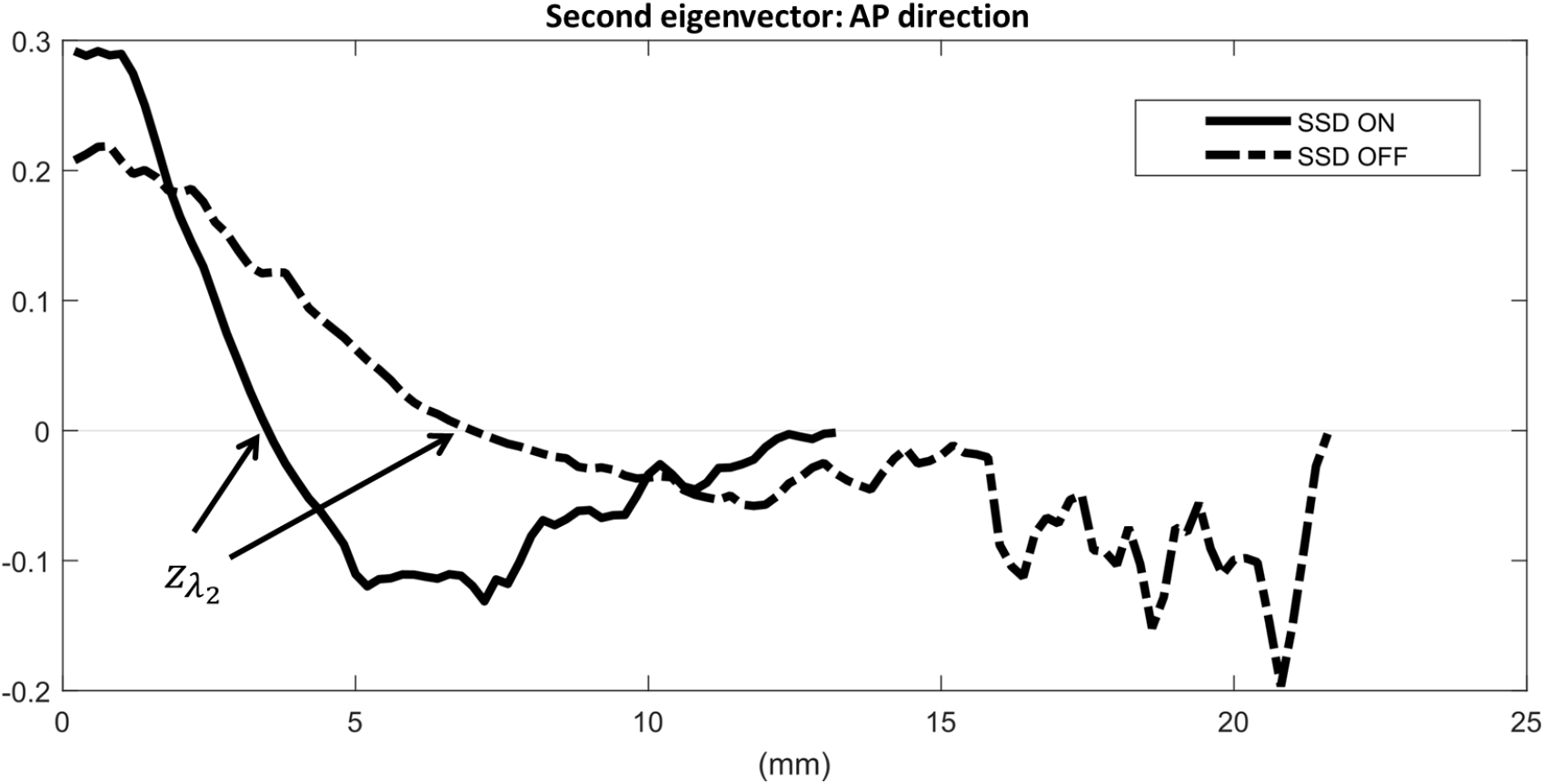
Representative plots of the second eigenvector of the transition matrix for both SSD ON (solid) and SSD OFF (dashed) conditions. The first zero crossing point is denoted 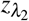. When additional skin stretch was applied on the subject (SSD ON), 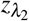 decreased. The scale of *y*-axis has no meaning, in contrast to the first eigenvector which is a probability measure. Instead, the sum of all components of the second eigenvector becomes zero.

IDA gives a comprehensive view on postural sway behavior and its long-term prediction. Table 1 shows that *D*_95,*Rad*_ tended to decrease (*p* = 0.07) when SSD was provided, suggesting that radial COP stayed closer to the centroid. A lower entropy (*H*_*AP*_, *p* = 0.05; *H*_*Rad*_, *p* = 0.04) with SSD ON implies a more deterministic system that one can more actively control the ankle and keeps their COP closer to a relative equilibrium position. It can be seen in the representative invariant density plot (Fig. 3) that *P*_*peak,AP*_ is greater (*p* = 0.05) and the COP location is more concentrated with SSD ON, compared with SSD OFF. It indicated that COP traveled around the centroid in a more regular pattern and it has a higher probability to be found in particular states (i.e., more likely to be closer to the centroid)^23^.

Among SDA measures (Table 1), both *D*_*s,ML*_ (*t*(14) = −2.42, *d* =0.62, *p* = 0.03) and ⟨Δ*r*^2^⟩_*c,Rad*_ (*t*(14) = −2.39, *d* = 0.62, *p* = 0.03) significantly decreased by approximately 15%. ⟨Δ*r*^2^⟩_*c,AP*_ tended to reduce when SSD was provided (*p* = 0.08). No significant effects were observed for the other SDA parameters. Figure 5 illustrates the representative stabilogram-diffusion plots from a subject for both SSD ON and OFF conditions. The critical point ⟨Δ*r*^2^⟩_*c*_ decreased in AP direction with a tendency (*t*(14) = −1.90, *d* = 0.49, *p* = 0.08) and in Radial direction with significance (*p* = 0.03) when the skin stretch feedback was provided. Looking at the representative linear stabilogram diffusion plot (Fig. 5), it can be inferred that skin stretch feedback reduced the rate of COP diffusion from the equilibrium point^20^. This also provides insight into how the skin stretch feedback enhances the open-loop and closed-loop control mechanisms on postural stability and demonstrates a reduced random behavior of COP.

**Figure 5.**
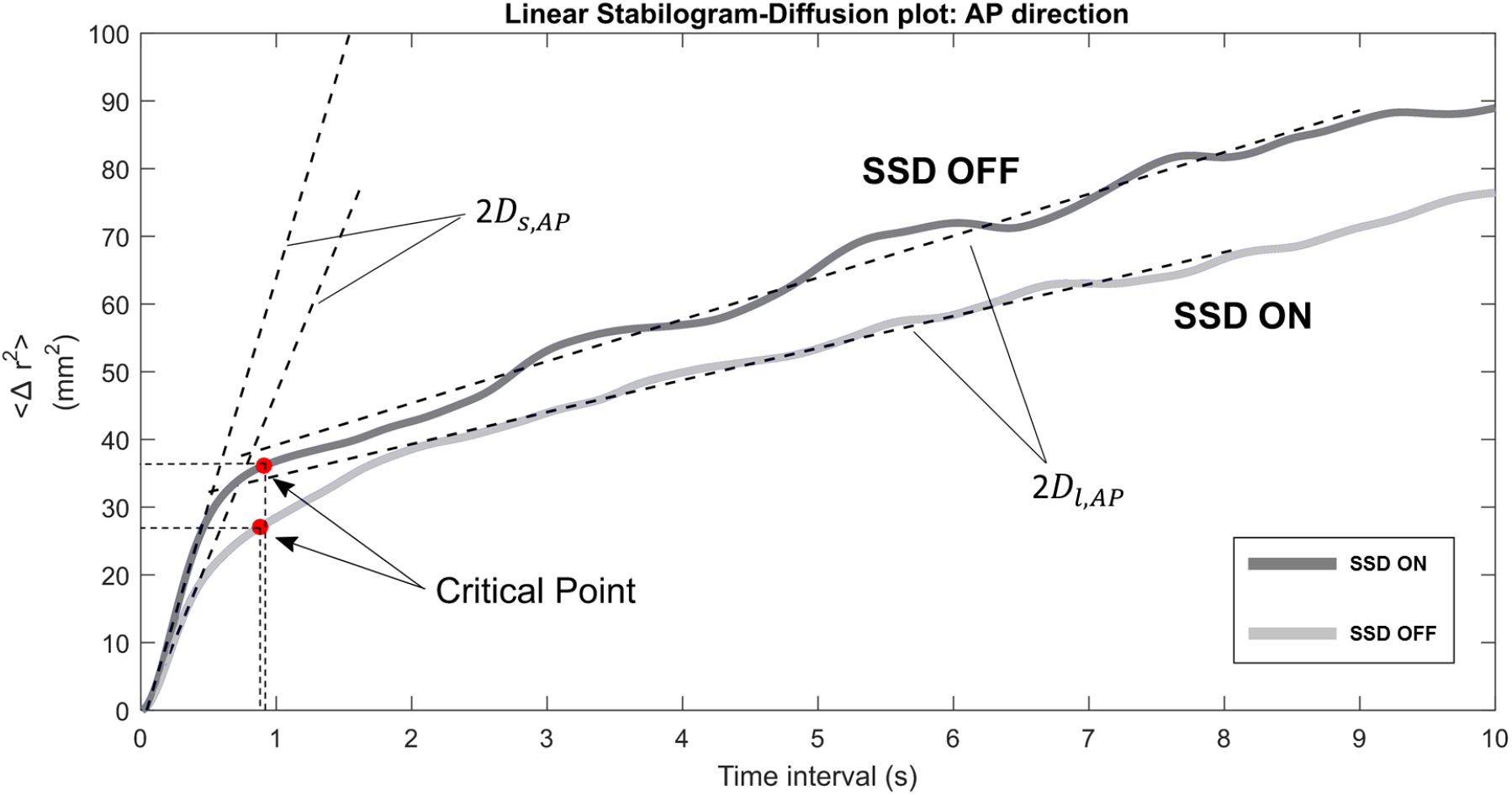
Representative stabilogram-diffusion plots from a subject. It shows two SDA plots of both SDD ON and SDD OFF conditions. The slopes of lines fitted to both short-term region (*D*_*s,AP*_) and long-term region (*D*_*l,AP*_) are less steep while SSD was on. The critical point coordinates (Δ*t*_*c,AP*_, ⟨Δ*r*^2^⟩_*c,AP*_) are both smaller when SSD ON.

## Discussion

In this paper, entropy and skin stretch feedback were examined based on the neuromechanical first principles and were considered for the design of the robotic balance rehabilitation system. The effectiveness of this system was examined in healthy young adults standing quietly with the perturbed sensory systems (i.e., removed vision and unreliable vestibular systems). IDA and SDA were performed to examine the efficacy of SSD during quiet standing in terms of stochastic processes.

The findings from IDA and SDA indicate that the postural sway significantly enhances when the additional sensory augmentation from skin stretch feedback is applied. The additional fingertip cutaneous sensation may have provided subjects with additional body sway information along the AP direction. Similar to the previous studies using the concept of light touch^32–34, 41, 49–51^, the fingertip cutaneous sensation replicated the sensory cues obtained from the fixed surface by a rotational contactor. However, unlike the previous studies that required a reachable stable surface, our SSD can be easily worn and carried by people, suggesting that it is a highly portable balance rehabilitation system.

According to the free-energy principle, the human postural control system adjusts the body posture to minimize the free energy of the system, or equivalently, to maximize the self-evidence based on the internal model^27^. The skin stretch feedback is an additional and artificial afferent signal that is fused in the sensorimotor system. By nature, existing afferent signals (e.g., proprioception, vestibular sensation, vision, somatosensation) are noisy and have uncertainty. Therefore, similar to Kalman filter, when the skin stretch feedback is combined with the existing afferent signals, the quality of the state estimation for the sensorimotor integration (e.g., body sway position, velocity and acceleration) improves^46^. Thus, the postural control system can easily increase the self-evidence of the system based on the internal model, which in turn, reduces the entropy.

The enhanced postural sway from IDA and SDA analyses due to the skin stretch feedback seems to be related to the strategy of the ankle stiffening. In other words, skin stretch feedback increased the ankle stiffness of the inverted pendulum model^7, 58, 59^. This agrees with previous studies regarding the influences of secondary tasks (e.g., internal/external focus on touching^60^, articulation task^61^) on postural control and suggests that the adaptive mechanism ensures a more active (increased frequency of sway) and more robust (reduced sway) control strategy. The present study confirms the theoretical findings in^62^ that if CNS receives sufficient body sway velocity information, ankle muscle activation can be modulated in anticipation of the changes in the COM position and therefore stabilizes the body, which reinforces the idea of free energy principle for postural control. It is possible that subjects used hip and knee strategies to reduce the entropy. However, it was visually confirmed that all subjects did not use any excessive movement of hip or knee. Also, due to the nature of the unperturbed quiet standing, it is believed that subjects used ankle strategy.

In this study, body sway velocity information was used to synthesize the controller of the skin stretch device rather than body sway position information^13^. In addition to position information, velocity information is essential in contributing to the regulation of posture balance^62^. Pan et al.^13^ reported that position-based skin stretch feedback did not have consistent positive effects on postural sway in various sensory deficit conditions. In the current study, we found that velocity-based skin stretch feedback significantly reduced the entropy of the postural sway. This may suggest that subjects learn the mapping between skin stretch feedback and body sway movements more quickly and efficiently by the velocity-based coding scheme rather than the position-based coding scheme. Sketch et al.^63^ found that a control scheme using velocity-based haptic feedback is more intuitive and effective compared with a control scheme using position-based haptic feedback. Bark et al.^64^ reported that subjects seem to have better ability in detecting changes in skin stretch stimuli than in the magnitude of the stimuli. This may support that subjects are more sensitive to body tilt rate instead of the absolute tilt angle. In sum, velocity-based skin stretch feedback may be more effective in minimizing the entropy of the postural sway. However, a further systematic investigation is required to confirm that this speculation.

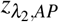 significantly decreased (*p* = 0.02) when skin stretch feedback was provided. Similarly, 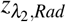 tended to decrease (*p*= 0.07) when skin stretch feedback was provided. As suggested by^55–57^, 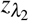 may represent the location in the state space where the dynamic behavior changes. Collins and De Luca^20^ observed two distinct regions of behavior during quiet standing and postulated that there exist both open loop and closed loop control regimes during quiet standing. This may indicate that the set of state 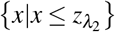 is the open loop control regime and the set of state 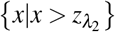 is the closed loop control regime. Following the interpretation of the SDA, postural sway randomly diffuses in the open loop region near the centroid. When the COP enters into the closed loop region (i.e., farther away from the centroid) due to external perturbations and the inherent instability of the upright standing, the postural control system actively adjusts the standing posture so that the postural sway enters back into the open loop region. Therefore, 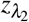 may suggest that the open loop region shrinks as the skin stretch feedback. is provided. It is very inspiring to note 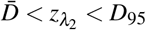. This inequality also supports the idea that 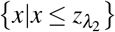 is related to the open loop control regime and 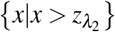is related to the closed loop control regime. However, it is not still known that 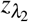 is the unique point where the control scheme changes. In other words, it is not sure whether the transition happens at a single point or in an overlapping area between the two control regimes. Further simulation-based research needs to verify this.

Interestingly, *D*_*s,ML*_ significantly decreased when skin stretch feedback based on postural sway in AP direction was provided, suggesting that skin stretch feedback information in AP direction significantly enhanced the open-loop control of posture in the ML direction. Since the skin stretch feedback contains the information of only AP directional postural sway, the enhancement of *D*_*s,ML*_ may seem nonsensical. Several studies have reported that postural sway is more sensitive in ML direction^16, 65, 66^. However, increased sensitivity in ML direction is not sufficient enough to explain this phenomenon. One possible explanation is that the skin stretch feedback may have worked as stochastic resonance. *Stochastic resonance* means that white noise signals can enhance the sensation of weak signals at some sensory receptors^67^. For example, a subthreshold random vibration at the hand could enhance the touch sensation at the fingertip of the stroke survivors^36^. Similarly, a subthreshold random vibration at the hand could enhance the motor performance of both healthy people and stroke survivors^37, 38^. Thus, if skin stretch feedback based on postural sway in the AP direction worked as a stochastic resonance in all directions, *D*_*s,ML*_ could enhance with the skin stretch feedback based on postural sway in AP direction. However, there should be one thing to be assumed for this to happen. Since skin stretch stimulus happens at the fingertip and other afferent signals are coming from different locations (e.g., ankle joint, foot, inner ear), the sensory fusion should happen at the supraspinal level. Luckily, Pan et al.^13^ showed the possibility that skin stretch feedback for the standing balance is happening at the supraspinal level. However, further systematic studies are needed to investigate the effect of random skin stretch feedback at the fingertip on the entropy of the postural sway.

This paper is not without limitations. The limb dominance of each subject was not considered. It was assumed that desensitization would not occur if moderate breaks were given between trials^68^, but this was not explicitly verified in this experiment. Adaptation to the device over the course of the trials was not examined. It is possible that the SSD may have not stimulated skin stretch only, but also have stimulated additional sensory modalities including vibration and pressure. It is important to note that these modalities are coupled: without pressure, there is no shear friction between the fingertip and SSD and thus no skin stretch. Also, shear friction can induce mechanical vibration due to elastic properties of fingertip. However, it is important to note that the skin stretch feedback due to shear friction at the fingertip may have provided the additional information of direction and speed of postural away even when there may have been small and/or consistent pressure and vibration.

## Conclusion

In this paper, we devised a way of quantifying the standing balance using a neuromechanical first principle of the sensorimotor system, called free-energy principle. Specifically, entropy was defined, and an algorithm for computing entropy of the human postural control system was investigated. Skin stretch feedback was examined, and a robotic balance rehabilitation system was designed so that the entropy of the postural control system reduces with the sensory augmentation (i.e., skin stretch feedback) according to the free-energy principle. An experiment was conducted and found that the proposed skin stretch device could effectively reduce the entropy of the postural control system when the proposed skin stretch device was used. Thus, we implicitly showed the feasibility of the framework of free energy principle in the human postural control system.

## Acknowledgements

This study was partially supported by fundings from National Institute for Occupational and Environmental Health (NIOSH)/Center for Disease Control and Prevention Education and Research Center at the Southwest Center for Occupational and Environmental Health (SWCOEH) (Grant #5T42OH008421) and the National Science Foundation Engineering Research Center for Precise Advanced Technologies and Health Systems for Underserved Populations (PATHS-UP) (Award #1648451).

## Author contributions statement

P.H. conceived and set up the experiment, analyzed the results, and wrote the manuscript. Y.P. conducted the experiment, analyzed the results and helped P.H. write the manuscript. C.D. analyzed the results and helped P.H. write the manuscript. All authors reviewed the manuscript.

## Data availability

The datasets generated and analyzed during the current study are available from github repository. https://github.com/pilwonhur/SensoryAugmentationVelocity/

## Appendix

The following shows that minimizing the free energy of the self-organizing biological system is implicitly equivalent to minimizing the surprise and entropy. Let *S* be the set of the sensory states, and *R* be the set of the internal states. For *s* ∈ *S*, *μ* ∈ *R*, and *m* a generative model, or forward model, a surprise is defined as (−log *p*(*s|m*)) which is a self-information and describes the accuracy of the model^69^. Kullback-Leibler divergence term appears in the free energy to resolve the intractability of the marginalization in computing the posterior density by introducing variational (or, approximate) density. Then, we find that the free energy is the upper bound of the surprise which is the log of negative self-evidence. In other words, the surprise is the lower bound of the free energy.

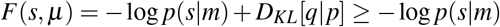

where *D*_*KL*_[·|·] is the Kullback-Leibler divergence between the posterior density and the variational density and is always nonnegative. Kullback-Leibler divergence becomes zero when the two densities are identical.

The transition matrix, *P*, of the postural sway is irreducible by the construction^23^ and aperiodic by its nature^70^. Note that the transition matrix, *P*, is an approximation of the Perron-Frobenius operator. Thus, *P* has the unique invariant density *π* and the process of the postural control system can be assumed to be ergodic^23, 71, 72^. Therefore, the long-term average of the free energy is the upper bound of the entropy of sensory states as shown below.

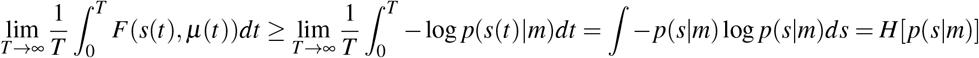

